# Interactions between motor thalamic field potentials and single unit spiking predict behavior in rats

**DOI:** 10.1101/642991

**Authors:** Matt Gaidica, Amy Hurst, Christopher Cyr, Daniel K. Leventhal

## Abstract

The thalamus plays a central role in generating circuit-level neural oscillations believed to coordinate brain activity over large spatiotemporal scales. Such thalamic influences are well-documented for sleep rhythms and in sensory systems, but the relationship between thalamic activity, motor circuit local field potential (LFP) oscillations, and behavior is unknown. We recorded wideband motor thalamic (Mthal) electrophysiology as healthy rats performed a two-alternative forced choice task. The power of delta (1*−*4 Hz), beta (13*−*30 Hz), low gamma (30*−*70 Hz), and high gamma (70*−*200 Hz) oscillations were strongly modulated by task performance. As in cortex, delta phase predicted beta/low gamma power and reaction time. Furthermore, delta phase differentially predicted spike timing in functionally distinct populations of Mthal neurons, which also predicted task performance and beta power. These complex relationships suggest mechanisms for commonly observed LFP-LFP and spike-LFP interactions, as well as subcortical influences on motor output.

## Introduction

Local field potential (LFP) oscillations are rhythmic fluctuations in the extracellular potential that emerge from, and may also regulate (Anastassiou et al., 2010), neuronal dynamics over a large spatiotemporal scale (Fries, 2015). Various aspects of the LFP including phase, amplitude and frequency are correlated with sensorimotor phenomena (Friston et al., 2015; Armstrong et al., 2018; Pesaran et al., 2018). Delta band (∼1-4 Hz) oscillations predict movement kinematics (Bansal et al., 2011), reaction time (RT, Lakatos et al., 2008; Stefanics et al., 2010; Hamel-Thibault et al., 2018), and sensory thresholds (Schroeder and Lakatos, 2009; Fiebelkorn et al., 2013). Beta oscillations (∼13*−*30 Hz) in the cortex and basal ganglia are enhanced under a variety of conditions including pre-movement hold periods (Donoghue et al., 1998; Saleh et al., 2010), isometric contractions (Baker et al., 1997), post-movement “rebound” (Pfurtscheller et al., 1996; Feingold et al., 2015), and parkinsonism (Brown, 2006; Mallet et al., 2008; Ellens and Leventhal, 2013). Beta oscillations are also correlated with prolonged RTs (Leventhal et al., 2012; Khanna and Carmena, 2017; Shin et al., 2017; van Wijk, 2017 Torrecillos et al., 2018) and slowed movement (Pogosyan et al., 2009; Lofredi et al., 2019). Conversely, movement onset is (usually) associated with decreased beta and increased gamma (∼60*−*100 Hz) power (Feingold et al., 2015; Tan et al., 2019, but see Leventhal et al., 2012).

In addition to correlations with behavior, LFP oscillations exhibit complex spatiotemporal relationships with each other and single unit activity. LFP coherence is common between brain regions, providing a potential mechanism to coordinate inter-regional activity (Fries, 2015; Wang et al., 2019). For example, beta oscillations occur in bursts simultaneously throughout cortical-basal ganglia circuits, with network-wide single unit activity locked to beta phase (Leventhal et al., 2012). Oscillations of different frequencies are commonly coupled to each other, both within and between brain regions. The possibility of an oscillatory “hierarchy” (Lakatos et al., 2005; Canolty et al., 2007) stems in part from observations that delta phase predicts beta oscillation amplitude (Saleh et al., 2010; López-Azcárate et al., 2013; Arnal et al., 2015; Hamel-Thibault et al., 2018; Grabot et al., 2019), and beta phase predicts the amplitude of higher frequency oscillations (de Hemptinne et al., 2013; Meidahl et al., 2019). These complex correlation patterns provide rich information regarding neural mechanisms of behavior, but make it difficult to distinguish cause from effect.

The thalamus is a central hub in nearly all motor, sensory, and associative circuits, and therefore well-positioned to regulate circuit-wide neuronal oscillations. Indeed, thalamocortical circuits generate or modulate many well-described LFP oscillations including sleep spindles (Halassa et al., 2011; Mak-McCully et al., 2017), cortical slow (< 1 Hz) oscillations (Neske, 2015), delta rhythms (Fogerson and Huguenard, 2016), alpha/mu (∼8*−*15 Hz) rhythms (Saalmann et al., 2012; Crunelli et al., 2018), beta rhythms (Bastos et al., 2014), and gamma rhythms (McAfee et al., 2018). Though many of these studies focused on sensory (especially visual) regions, motor thalamic (Mthal) spiking is also phase-locked to delta oscillations under anesthesia (Nakamura et al., 2014). Modeling studies suggest that motor system beta oscillations could result from layer-specific thalamocortical inputs (Sherman et al., 2016; Reis et al., 2019), though mechanisms intrinsic to the basal ganglia have also been proposed as “beta generators” (McCarthy et al., 2011; Tachibana et al., 2011; Mirzaei et al., 2017). Given the strong associations between thalamic activity and brain rhythms across sensory modalities and brain states, we hypothesized that Mthal, which is reciprocally connected with motor and premotor cortices, mediates many LFP-LFP and LFP-behavior correlations.

To understand the relationship between Mthal spiking, Mthal LFPs, and behavior, we recorded wideband Mthal activity as rats performed a two-alternative forced choice task. Using this data set, we previously found that two distinct, functionally defined populations of Mthal neurons predict dissociable aspects of task performance (Gaidica et al., 2018). Here, we describe Mthal LFP-behavior correlations, as well as novel relationships between functionally defined Mthal single units and LFP oscillations. These results have important implications for models of motor system LFP generation, as well as their functional interpretation.

## Results

### LFP Power and Phase are Modulated by Task Performance in Discrete Frequency Bands

Rats (n = 5) were cued to immediately move left or right from a center nose port based on the pitch of an instructional cue (Figure 1, “Tone” event) until a high degree of accuracy was achieved (77 ± 17% over 30 sessions, mean ± s.d.). Reaction times (RT; the time from Tone to Center Out) and movement times (MT; the time from Center Out to Side In) were consistent with similar studies (197 ± 10.3 ms and 302 ± 127 ms, respectively, mean ± s.d., see Gaidica et al, 2018 for the full distributions) (Dowd and Dunnett, 2005; Leventhal et al., 2012; Schmidt et al., 2013; Leventhal et al., 2014). Similar to observations in cortex (Murthy and Fetz, 1992; Saleh et al., 2010; Igarashi et al., 2013) and the basal ganglia (Berke et al., 2004; Masimore et al., 2005), the awake LFP power spectrum in Mthal had discrete peaks in delta (1–4 Hz), theta (4–7 Hz), beta (13–30 Hz), and low gamma (30–70 Hz) bands (Figure 1).

**Figure 1.**
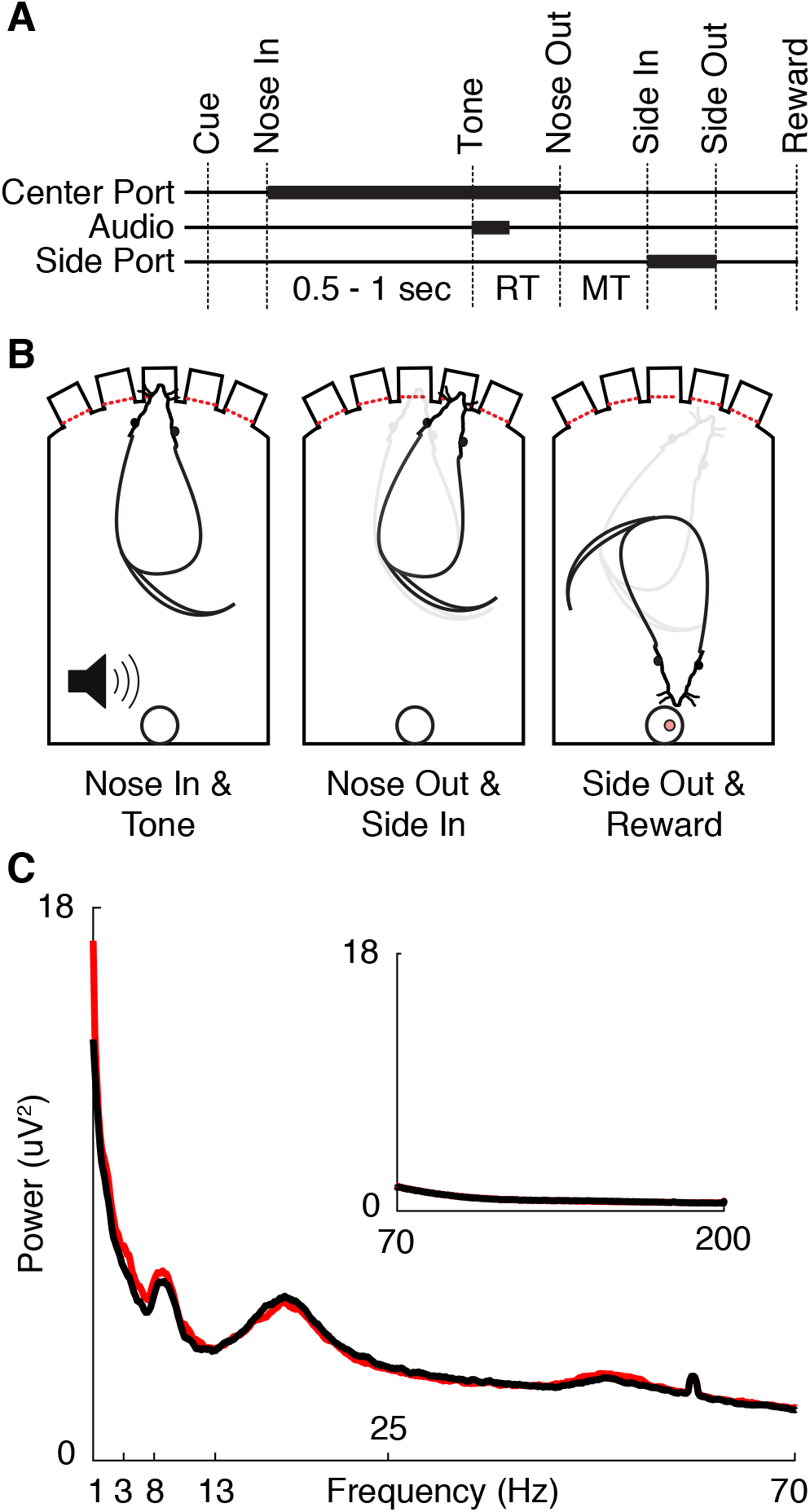
Behavioral task and physiology. **(A)** Trials began by illuminating one of the three center ports in a five-port behavior chamber (“Cue”). The rat poked and held its nose in the lit port (“Nose In”) for a variable interval (0.5–1 s, pulled from a uniform distribution) until a 1 or 4 kHz auditory cue (“Tone”) instructed the rat to move one port to the left or right, respectively. Nose Out, Side In, and Side Out indicate when the rat withdrew from the central port, poked the adjacent port, and withdrew from the adjacent port, respectively. “Reward” indicates the time of reward pellet retrieval. Reaction time (RT) and movement time (MT) intervals are labelled. Thick lines indicate either nose port occupancy (Center and Side Port) or playing the auditory cue (Audio) **(B)** Schematic of the rat operant chamber during key behavioral epochs. **(C)** Session-averaged power spectrum of low (1–70 Hz) and high (inset, 70–200 Hz) frequencies for in-trial (black) and inter-trial (red) periods.

Task-linked Mthal LFP power modulation was nearly identical to prior observations in motor cortex and the basal ganglia during a similar task (Figure 2) (Leventhal et al., 2012). Beta/low gamma power concurrently and transiently increased near the Nose Out and Side Out events, contradicting the widely-held view that beta power decreases with movement onset. This is likely explained by the delivery of instructive and imperative signals with the same stimulus - when these cues are separated, beta power increases during the inter-stimulus “hold” period and decreases with movement onset (including in our own experiments) (Donoghue et al., 1998; Saleh et al., 2010; Leventhal et al., 2012; Khanna and Carmena, 2017). Delta power also increased at Nose Out, and remained elevated through Side In. Finally, high gamma power transiently increased at Nose-and Side Out, and exhibited a sustained elevation as the rat moved from the nose ports to the food receptacle (prior to the Reward event).

**Figure 2.**
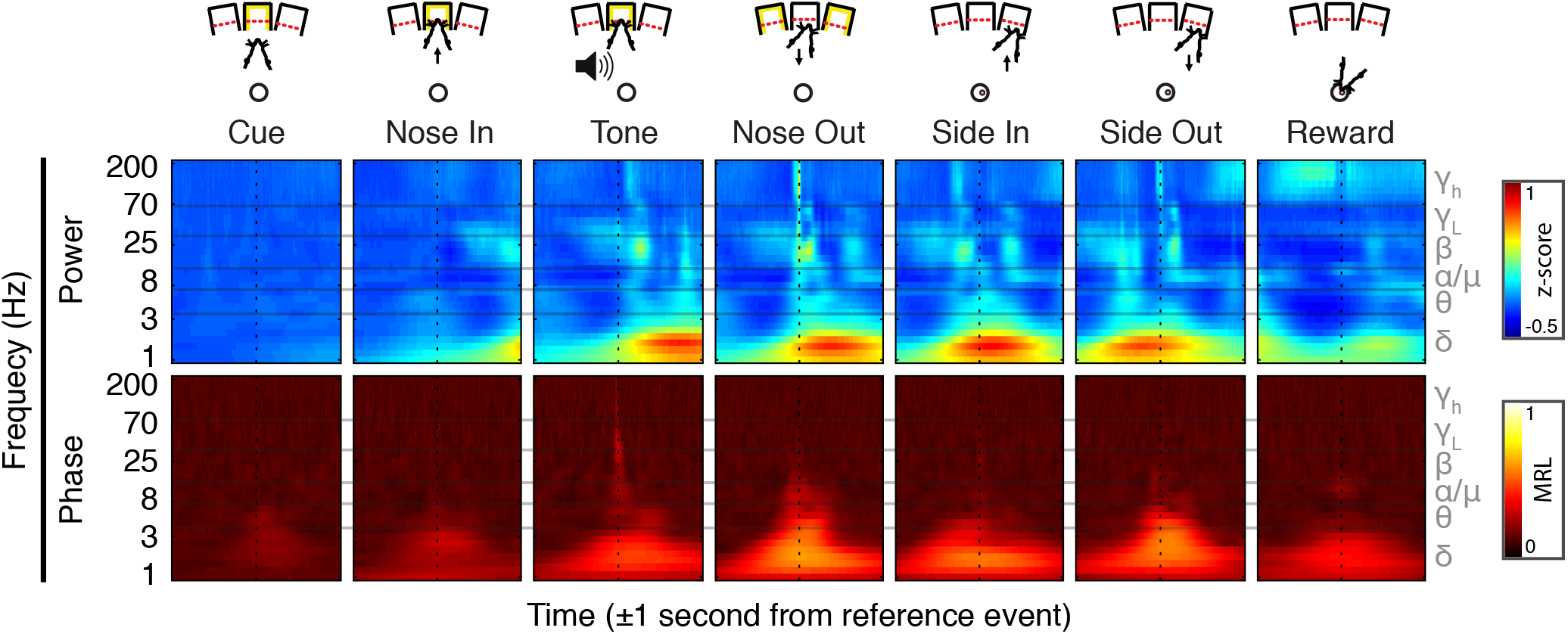
Peri-event LFP power and phase modulation. (Top) behavioral schematic for a rightward-cued successful trial. (Middle) Mean gabor spectrograms for each event. (Bottom) Mean resultant length (MRL) of event-locked LFP phase. Higher values indicate time-frequency points at which LFP phase tends to be aligned across trials. The data used to generate Figure 2 and associated Matlab code are included in Figure 2 – Source Data 1.

To determine if the comodulation of beta and low gamma power results from independent modulation of these frequency bands by the same behavioral events, we performed two additional analyses. First, correlations between beta and low gamma power were also present, albeit weaker, when the rats were not actively engaged in the task. Second, these correlations nearly disappeared when recalculated using trial-shuffled data (Figure 3). These findings argue that beta/low gamma power-power correlations are a general feature of Mthal physiology.

**Figure 3.**
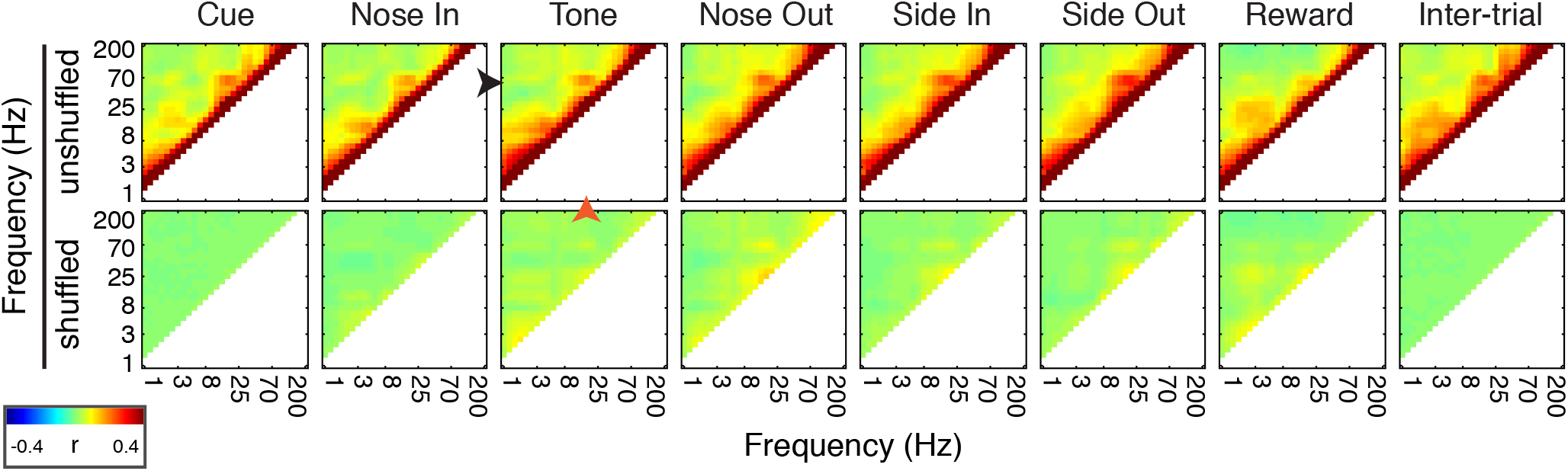
Mthal LFP power in discrete frequency bands is comodulated during and between trials. (Top) power-power comodugrams locked to each behavioral event and during the inter-trial interval. Note the consistent positive correlation between continuous beta (20 Hz, orange arrowhead at Tone) and low gamma (55 Hz, black arrowhead at Tone) power. (Bottom) comodugrams for the same events calculated using trial-shuffled data. The data used to generate Figure 3 and associated Matlab code are included in Figure 3 – Source Data 1.

In addition to LFP power changes, LFP phase in specific bands was strongly modulated by the task. Beta/low gamma phase became sharply aligned at the Tone event (Figure 2), as previously observed in the basal ganglia (Leventhal et al., 2012). Phase alignment in the delta band was present as early as the Cue event and peaked at the Nose Out and Side Out events.

Collectively, these data suggest complex temporal coordination of LFP power and phase in discrete frequency bands.

### Delta Phase Predicts Beta and Low Gamma Power

The co-occurrence of a delta phase alignment and beta/low gamma power increase at Nose Out suggests that phase-amplitude coupling (PAC) is a prominent feature of Mthal physiology, as has been observed in other brain regions (Canolty et al., 2006; Tort et al., 2008; Cohen et al., 2009; Dejean et al., 2011; Belluscio et al., 2012; López-Azcárate et al., 2013). Indeed, delta-beta/low gamma PAC was significantly elevated throughout the task (“in-trial”, Figure 4), most prominently during movement from the Center to Side nose ports (i.e., Nose Out to Side In, Figure 4). Significant delta-beta/low gamma PAC was also present when the rat was not actively engaged in the task (“inter-trial”), and was significantly diminished when recalculated using trial-shuffled data. As for beta/low gamma amplitude-amplitude coupling, these results argue that delta-beta/low gamma PAC does not result simply from common responses to behavioral events.

**Figure 4.**
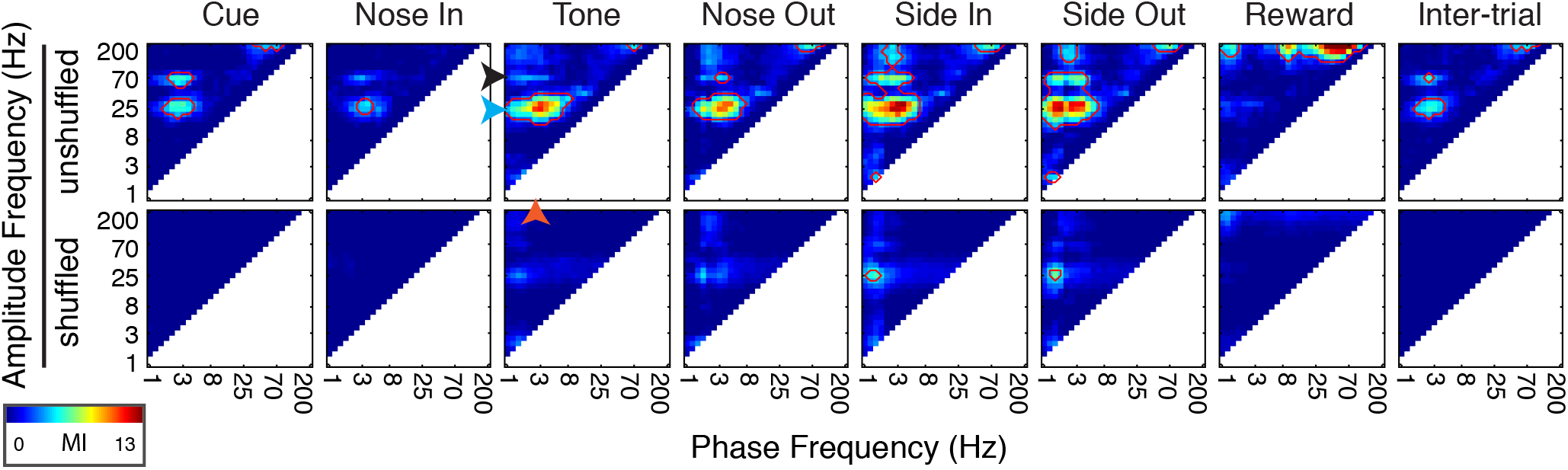
Phase-amplitude coupling (PAC) is dynamically modulated by task events. (Top) Peri-event PAC as assessed by the modulation index (MI, see Materials and Methods). Arrows indicate specific frequencies in the delta (2.5 Hz, orange arrowhead), beta (20 Hz, blue arrowhead), and low gamma (55 Hz, black arrowhead) bands. (Bottom) same calculation using trial shuffled data. Red outlines highlight areas where PAC is significant (p < 0.05). The data used to generate Figure 4 and associated Matlab code are included in Figure 4 – Source Data 1.

### Delta Phase Predicts Single Unit Mthal Activity

The phase of low frequency oscillations also predicted the timing of single unit activity (Lakatos et al., 2005 Fujisawa and Buzsáki, 2011; Nakamura et al., 2014; Crunelli et al., 2015). 59% of all units (n = 366) exhibited a non-uniform delta phase distribution during trials (black line in Figure 5; defined as p < 0.05 for each unit, Rayleigh test for non-uniformity), which fell to 36% during the inter-trial period. These percentages were significantly greater than chance, as assessed by surrogate firing-rate matched Poisson spike trains. Furthermore, spike-phase entrainment was unique to the delta and theta bands. The average mean resultant length (MRL, a measure of phase uniformity) of spike-LFP phases across units was also significantly greater than chance for low frequencies (p < 0.001 at 2.5 Hz). These data support the notion that low-frequency oscillations modulate Mthal single neuron excitability in a behaviorally-relevant manner.

**Figure 5.**
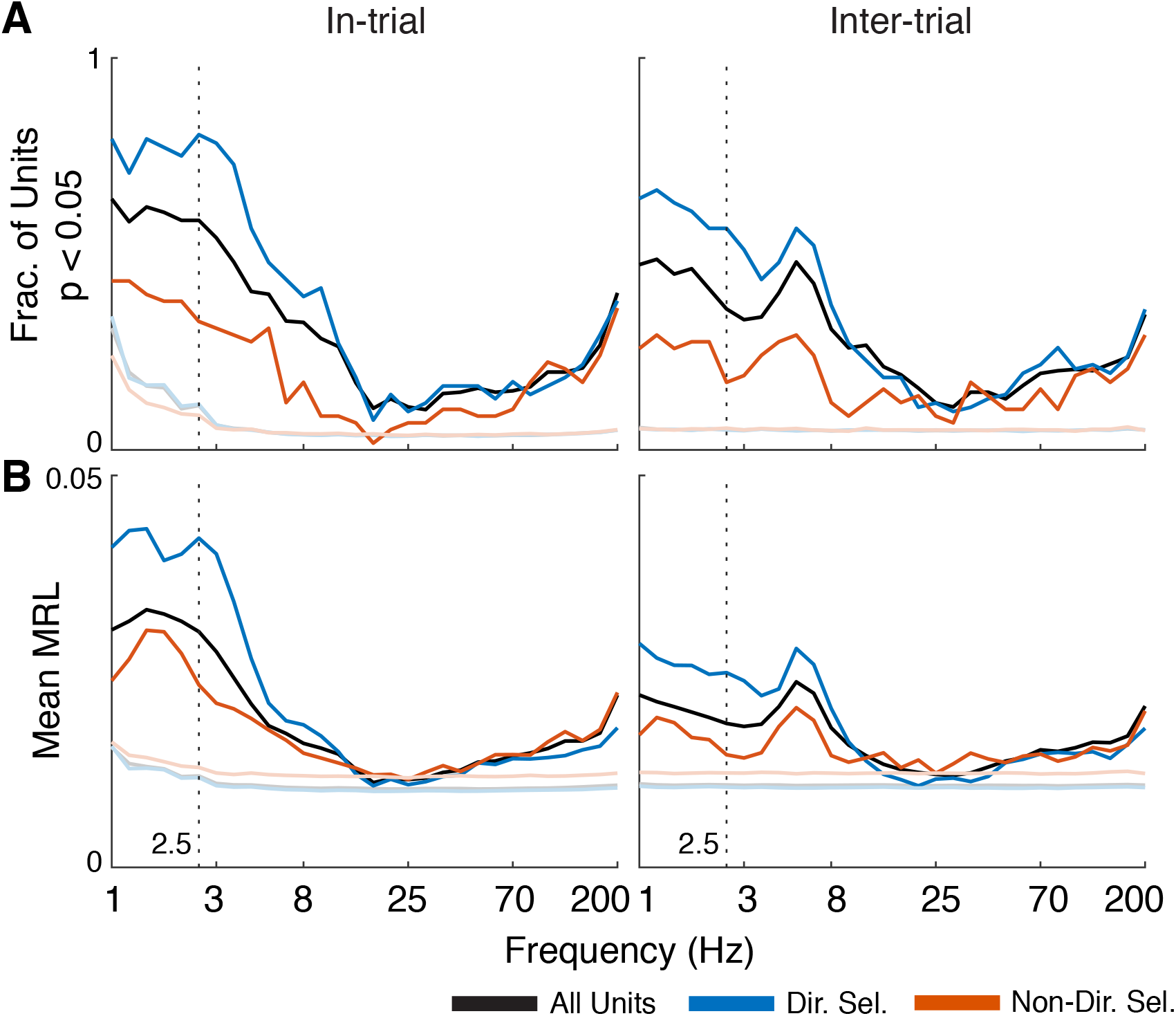
Single unit activity is selectively entrained to low frequency oscillations. **(A)** The fraction of units from each population (all units, directionally selective units, and non-directionally selective units) that were significantly entrained to LFP oscillations across frequencies (p < 0.05, Rayleigh test for non-uniformity) during task engagement (“in-trial”) and during the inter-trial interval (“inter-trial”). The fraction of units entrained at 2.5 Hz was significantly different from the Poisson-distributed surrogate spike trains for each unit population during both In-Trial and Inter-Trial epochs (p < 0.001) **(B)** Average mean resultant length (MRL) for each unit population across frequencies. The mean MRL for each population was significantly different from the surrogate spike trains at 2.5 Hz (p < 0.001). The data used to generate Figure 5 and associated Matlab code are included in Figure 5 – Source Data 1.

We next investigated whether phase preferences differed for two functionally distinct subpopulations of Mthal units previously identified in this data set (Gaidica et al., 2018). Briefly, “directionally selective” unit activity was tightly linked to the Nose Out event, predicted which direction the rat would move, and predicted both RT and MT. Conversely, “non-directionally selective” units were more tightly locked to the Tone event and predicted RT, but not MT or movement direction (366 total units, 103 directionally selective units, and 75 non-directionally selective units).

These functionally defined populations were differentially entrained to delta oscillations. In-trial, 80% of directionally selective units were significantly entrained to delta phase (Figure 5), which was the case for only 33% of non-directionally selective units. Between trials, delta entrainment decreased slightly for all units, resulting in entrainment for 57% of directionally selective units and 17% of non-directionally selective units. At higher (alpha/beta) frequencies, the entrainment for all three groups (directionally-selective, non-directionally selective, and all units) approached chance as assessed by surrogate Poisson spike trains. Thus, most of the single unit delta entrainment was due to directionally-selective units, and this entrainment was specific to low frequencies.

To determine if Mthal units tended to fire at the same preferred delta phase (assessed at 2.5 Hz), we created a spike-phase histogram for each unit (Figure 6). Within trials, there was a clear phase preference for both directionally and non-directionally selective units (175.69°, p = 5.7 *×* 10^-7^ and 113.86°, p = 0.0044, respectively, Rayleigh test for non-uniformity). Between trials, the phase preference for non-directionally selective units disappeared. For directionally selective units, however, the phase preference persisted (191.13°, p = 9.5 *×* 10^-8^ Raleigh test for non-uniformity) and was statistically indistinguishable from the in-trial phase preference (p = 1 compared with in-trial phase, Kuiper two-sample test against the null hypothesis that the two distributions are identical). These results suggest that the in-trial phase entrainment observed for non-directionally selective units may be an artifact of two physiologic events independently locked to the same behavioral event (spiking and delta phase alignment). Conversely, the phase entrainment of directionally selective units is more likely a pervasive feature of Mthal physiology.

**Figure 6.**
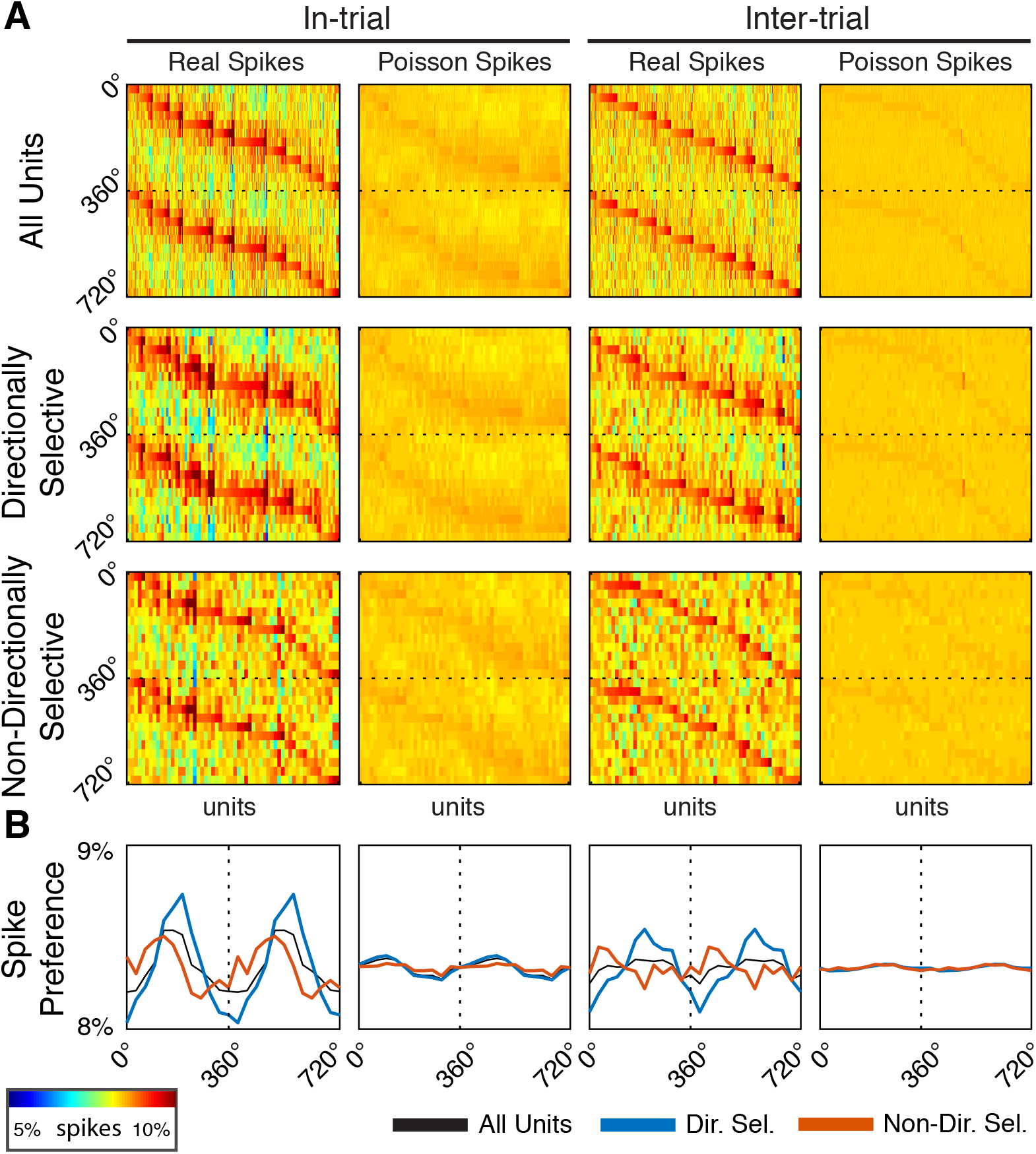
Single unit entrainment occurs at preferred delta phases specifically for directionally-selective units. **(A)** Spike-phase histograms for functionally-defined single unit populations and surrogate Poisson-distributed spike trains. Each column within the individual phase histograms represents a single unit. Colors indicate the percentage of spikes within a phase bin (12 bins from 0° to 360°, repeated to 720° for clarity). Units are sorted by their preferred phase separately for each plot. **(B)** Mean spike-phase histograms for each unit population. The data used to generate Figure 6 and associated Matlab code are included in Figure 6 – Source Data 1.

### Directionally Selective Unit Activity Uniquely Predicts LFP Power

Delta phase predicts both beta/low gamma power and single unit spiking. We therefore hypothesized that spiking and beta/low gamma power are also correlated. To test this, we cross correlated LFP power with a continuous spike density estimate (SDE) of all Mthal single unit activity and compared it to chance using a Poisson spike distribution.

During trials, directionally selective unit activity was maximally correlated with beta power (r = 0.03) at a lag of −0.72 s (i.e., Mthal spiking preceded beta power increases on average, Figures 7 and 8). There was a smaller, yet significant negative correlation (r = −0.02) that peaked at −0.4 s, which may reflect decreased Mthal activity preceding the Nose-and Side In events when beta power is enhanced (see Gaidica et al., 2018 Figure 2). The cross-correlation pattern was strikingly similar during inter-trial intervals but attenuated (r = 0.01 at −0.58 s lag, r = −0.014 at −0.25 s lag), suggesting that beta power is enhanced following a “pause-fire” pattern of directionally selective Mthal unit spiking (Figure 8). Non-directionally selective unit activity was correlated with beta power slightly earlier, and to a lesser degree in-trial (r = 0.016 at t = −0.18 s lag, r = −0.017 at t = −0.52 s lag), but was not significantly correlated with beta power during the inter-trial period. These results suggest that the relationship between non-directionally selective unit activity and beta power resulted from independent correlations with behavioral events. Conversely, the relationship between directionally selective unit activity and beta power is likely a general feature of Mthal physiology.

**Figure 7.**
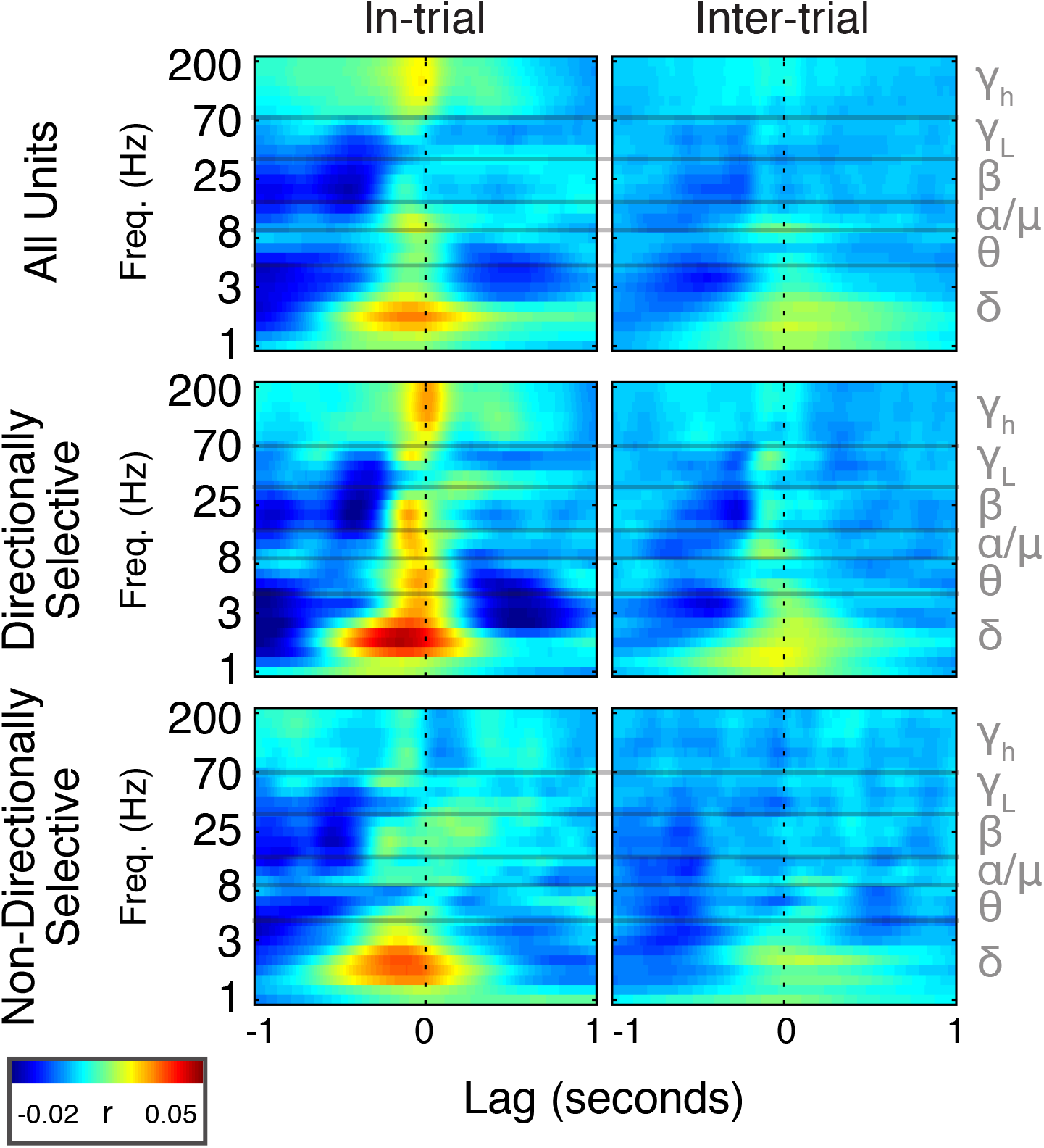
Spike timing is correlated with power modulation in specific frequency bands. **(A)** Each horizontal line indicates the average cross-correlation between individual unit spike density estimates and LFP power at different frequencies. t = 0 indicates a zero-lag correlation. Negative lags indicate that spikes lead LFP changes, and positive lags indicate that LFP changes lead spiking.

**Figure 8.**
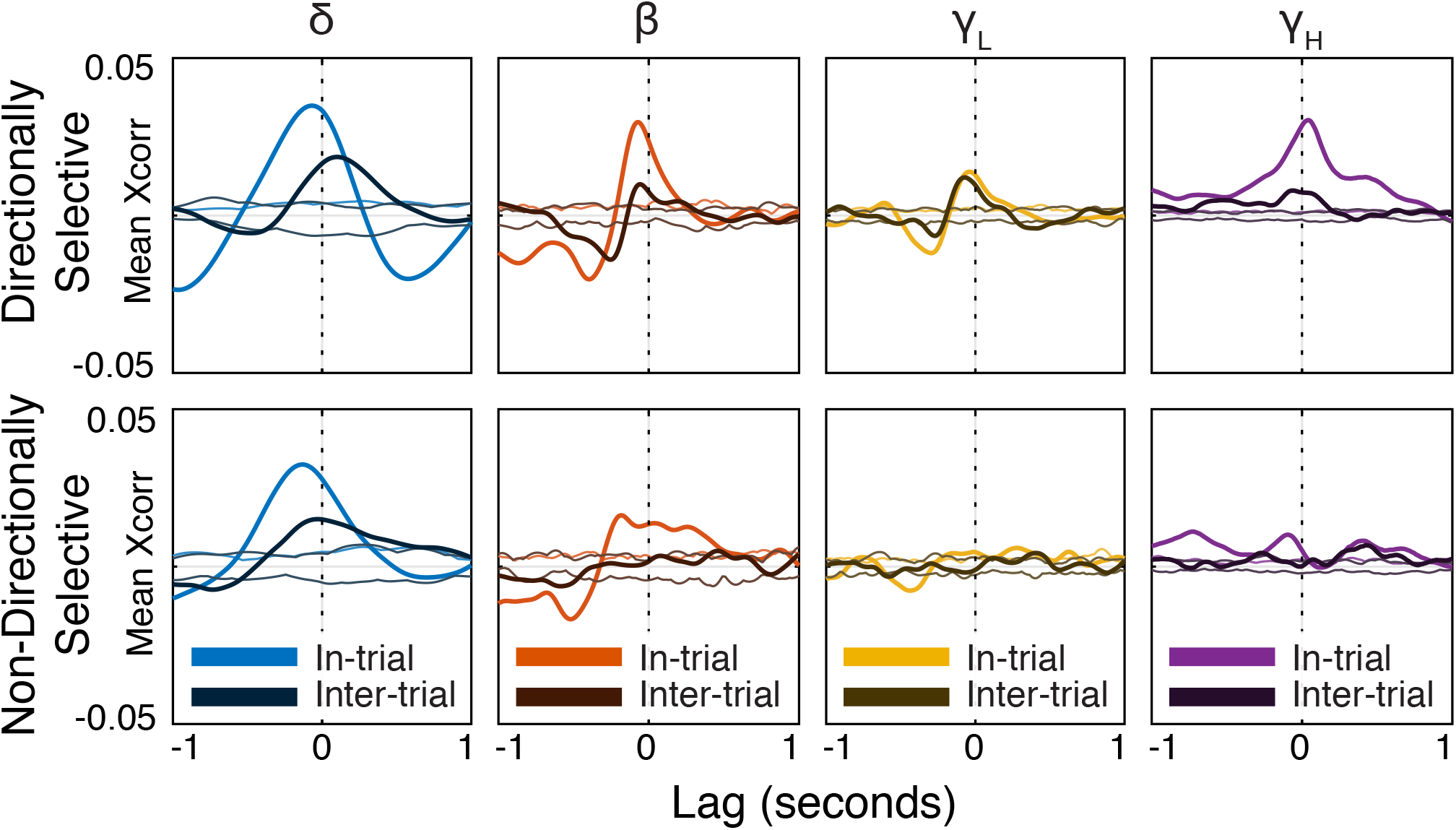
Beta power lags directionally-selective unit spiking during and between trials. **(A)** Frequency band-specific cross correlations (±1 s) for directionally selective (top) and non-directionally selective (bottom) units. Thick lines represent the actual cross-correlations; thin lines represent thresholds for significance (see Materials and Methods). Note that the cross-correlation patterns between directionally-selective units and beta/low gamma power are similar both during and between trials, which is not true for any other unit-frequency band combinations.

Similar patterns were observed for directionally selective unit spike-low gamma power correlations, which were significant during both in-trial and inter-trial epochs. However, non-directionally selective unit activity was not correlated with low gamma power during either epoch. The consistency of these correlations (or lack thereof) across both behavioral epochs supports the notion that directionally selective unit activity is uniquely linked to the LFP.

The pattern of high gamma modulation during the task closely matched single unit Mthal activity patterns (Gaidica et al., 2018, Figure 2), consistent with observations that high frequency oscillations are correlated with multi-unit activity. High gamma power best correlated with directionally selective unit activity, exhibiting roughly zero-lag between spiking and power increases. Therefore, as in cortex, Mthal high gamma power may serve as a surrogate for multi-unit activity (Ray et al., 2008; Manning et al., 2009; Watson et al., 2018).

Mthal single unit activity also showed a small correlation with delta power, which was larger for directionally selective units. Unlike the beta power correlation, the time lag and pattern of spike-delta power correlations was inconsistent between the in-trial and inter-trial periods (Figure 8). The peak spike-power correlation occurred at −0.68 s in-trial (r = 0.035) but at 0.1 s during the inter-trial interval (r = 0.019). Similar but smaller correlations were also observed for non-directionally selective units (r = 0.032 at −0.13 s in-trial, r = 0.015 at 0.04 inter-trial).

In summary, the consistency of in-trial and inter-trial correlations argues for a unique physiological relationship between directionally selective unit activity and beta/low gamma power in Mthal.

### LFP Correlates of Performance

Given the relationships between single unit activity and task performance (Gaidica et al., 2018), and single unit activity and LFP features, we next examined relationships between LFP features and task performance.

Delta phase near the Tone event strongly predicted RT (p < 0.05) in 19/30 recording sessions (Figure 9; session-averaged r = 0.42 at t = 0.53 s after the event). This suggests that there is a preferred Mthal delta phase for movement initiation (Figure 2), and that RT is (at least partially) determined by the distance from that preferred phase when the Tone plays (Lakatos et al., 2008). While we cannot completely rule out the possibility that filtering propagates a delta phase reset at Nose Out back in time to the Tone event (de Cheveigné and Nelken, 2019), similar delta phase-RT correlations have been reported in a range of behavioral paradigms (Stefanics et al., 2010 Hamel-Thibault et al., 2018 Saleh et al., 2010). Furthermore, while phase discontinuities were occasionally observed in the filtered signal (Figure 10A, orange marker), they were not consistently present at the Nose Out event (Figure 10C). Indeed, delta phase varied smoothly from the Tone through Nose Out events across all trials (Figure 10C).

**Figure 9.**
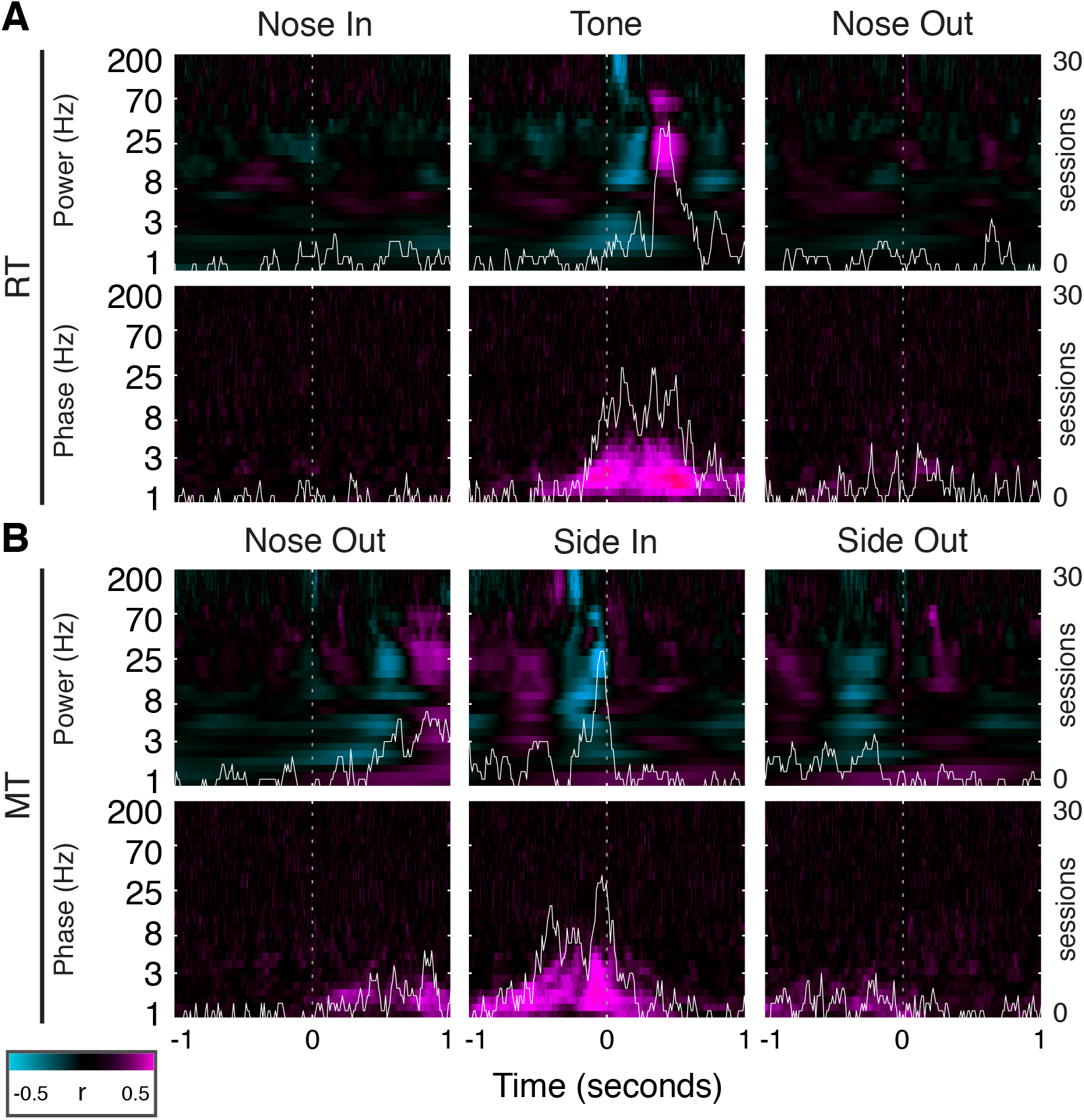
LFP oscillations predict task performance. **(A)** Peri-event (±1 s) reaction time (RT) correlations for power (top) and phase (bottom) for the Nose In, Tone, and Nose out events. Correlation values are session-averaged. The white lines indicate how many sessions reached significance (p < 0.05) at each time point in the beta band (20 Hz) for power, and in the delta band (2.5 Hz) for phase. **(B)** Same data as (A) for movement time (MT) from the Nose Out, Side In, and Side Out events. The data used to generate Figure 9 and associated Matlab code are included in Figure 9 – Source Data 1.

There was a similar delta phase correlation near the Side In event for MT (Figure 9B, p < 0.05 for 20/30 sessions; session-averaged r = 0.37 at t = 0.07 s before the event). However, since delta phase was aligned at Nose Out, and MT was approximately the length of a single delta oscillation cycle, one would expect Side In to occur at different delta phases for different MTs. Thus, this delta phase-MT correlation likely does not represent a new finding independent of the Nose Out delta phase alignment.

Beta power also predicted RT in the peri-Tone period (p < 0.05 for 21/30 sessions; session-averaged r = 0.29 at t = 0.45 s after the event) (Leventhal et al., 2012). As for the delta phase-MT correlation, however, this relationship can be explained by event-related beta modulation. Specifically, for short RT, beta power after the Tone event increases earlier because, by definition, the Nose Out event is closer to the Tone event. We previously reported a small but significant correlation between striatal beta power and RT in the immediate pre-Nose Out period (Leventhal et al., 2012), but this finding was not replicated in Mthal (p < 0.05 in only 4/30 sessions). Whether this is due to subtle differences between basal ganglia and Mthal physiology, failure to detect a subtle correlation in the present study, or a false positive result in the prior study, is unclear.

Finally, beta power was anticorrelated with MT just prior to Side In (p < 0.05 for 19/30 sessions; session-averaged r = −0.27 at t = 0.04 s before the event). However, because beta power increases transiently after Nose Out, beta power must be elevated just prior to Side In for short MT. This correlation is also, therefore, unlikely to represent a new effect independent of task-linked beta modulation. In summary, delta phase at the Tone event was the only LFP feature that consistently and independently predicted task performance.

## Discussion

We identified several interrelated correlations between Mthal LFPs, Mthal single unit activity, and behavior. First, LFP phase in the delta band, and power in multiple frequency bands (delta, beta, low and high gamma) were modulated by specific behavioral events. Delta phase strongly predicted RT, LFP beta/low gamma power, and single unit spike timing. Given these correlations, it is not surprising that spike timing also predicted beta/low gamma power, though we did not find an independent relationship between beta power and RT. Interestingly, Mthal single unit subpopulations previously identified on the basis of behavioral correlations (Gaidica et al., 2018) exhibited distinct relationships with delta phase and beta power. Many of these correlations persisted during the intertrial interval, arguing that they do not arise simply from independent correlations with behavior. These observations unify prior observations of correlations between delta phase, beta power, and behavior. They also provide new insights into how motor system LFP oscillations may be generated and linked to behavior.

Beta oscillations are suggested to represent a stabilized network state during which motor plans are less likely to change (Gilbertson et al., 2005; Pogosyan et al., 2009; Engel and Fries, 2010; Khanna and Carmena, 2017), which may serve the adaptive purpose of preventing distractors from interfering with a recently adopted plan. This interpretation is supported by small, but significant and reproducible, correlations between beta power and RT (Leventhal et al., 2012; Khanna and Carmena, 2017; Shin et al., 2017; van Wijk, 2017; Torrecillos et al., 2018). However, we did not replicate that finding here. This could be due to differences in recording sites, as prior correlations were found in basal ganglia or cortex. However, patterns of event-related beta power modulation were nearly identical in striatum and Mthal (Leventhal et al., 2012), making it less likely that Mthal and cortical-basal ganglia beta oscillations differentially predict RT. We suggest instead that beta power is linked to RT indirectly via delta-beta PAC, explaining why weak beta-RT correlations are frequently observed.

**Figure 10 with 1 supplement.**
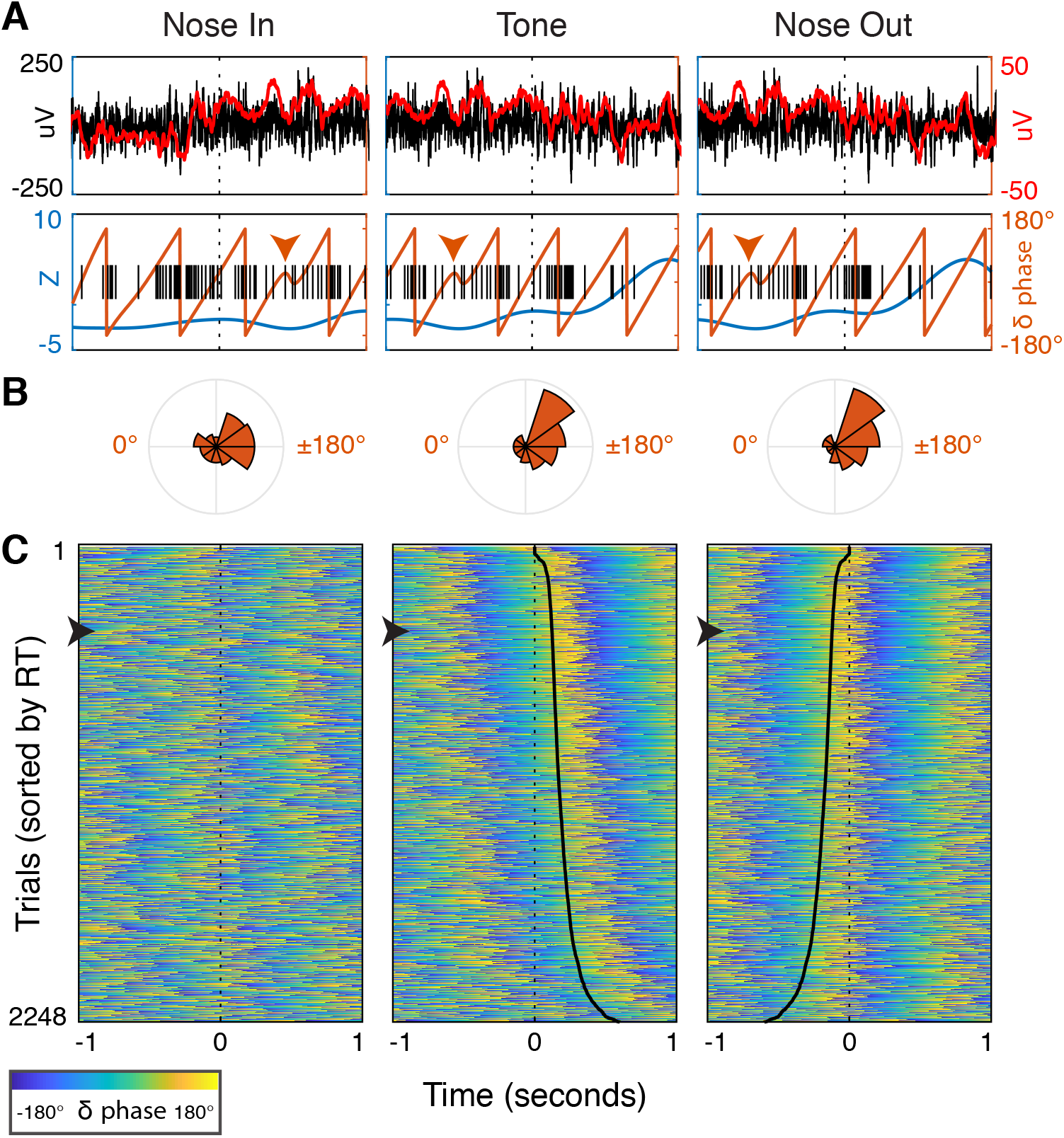
Delta phase predicts spiking and becomes aligned at Nose Out. **(A)** Peri-event (±1 s) data from a single trial. Top - the unfiltered, wideband signal (black, left axis) with a 50 ms smoothing filter (red) highlighting discrete delta-band oscillatory patterns after Nose In and prior to Tone. Bottom - delta power (blue) and phase (orange line, orange arrowhead highlights phase discontinuity), and single unit spike timing (black, directionally selective unit #319, bottom). **(B)** Normalized, single unit spike-phase histograms for the data in (A). **(C)** Peri-event delta phase from all trials sorted by RT. The trial from A is marked with a black arrowhead along left border. Black lines in the Tone and Nose Out panels indicate the Nose Out and Tone events, respectively (i.e., the time from reference event to the black line is RT).

Delta phase was more strongly and consistently correlated with RT prior to movement onset than beta power. Similar correlations have been found during tasks in which cortical delta oscillations entrain to rhythmic stimuli (Lakatos et al., 2008; Stefanics et al., 2010; Arnal et al., 2015). LFP oscillations may modulate neuronal excitability through ephaptic interactions (Anastassiou et al., 2010; Tiganj et al., 2014), or simply reflect aggregate synaptic drive that influences spiking probability (Pesaran et al., 2018). In either case, active entrainment of LFP oscillations to rhythmic cues is a potential mechanism to optimize neuronal excitability at the time of anticipated salient stimuli (Schroeder and Lakatos, 2009). It remains unclear, however, whether such mechanisms are generalizable to single interval timing (Breska and Deouell, 2017 Hamel-Thibault et al., 2018; Zoefel et al., 2018). In our task, instructive/imperative cue (Tone) timing is somewhat predictable, occurring 0.5–1.0 s after Nose In. The presence of significant delta phase coherence across trials even prior to Nose In (Figure 10–figure supplement 1), and the smooth progression of delta phase at Nose Out (as opposed to an abrupt phase reset, Figure 10), support the idea that delta phase actively aligns to increase the probability that the Tone arrives at a favorable phase for quick reactions.

A plausible mechanism for delta phase-RT correlations is that delta phase predicts (perhaps influences) Mthal spike timing, which drives motor cortex to initiate movement. Thus, if the Tone arrives just after the optimal phase, a full delta cycle would have to repeat before Mthal neurons are maximally excitable, suggesting a source of RT variability. In support of this hypothesis, units whose activity predicted RT were strongly entrained to delta rhythms. A related but slightly different interpretation is that circuit-wide delta oscillations simultaneously reflect thalamic and cortical excitability, with cortical neurons more likely to fire at specific delta phases independently of thalamic input (Lakatos et al., 2005; Rule et al., 2018).

Our data also suggest a mechanism for delta-beta PAC. Delta phase predicts Mthal single unit spike timing which in turn predicts, and possibly causes, cortical beta oscillations that are propagated throughout basal ganglia-thalamocortical circuits (Jones et al., 2009; Sherman et al., 2016; Reis et al., 2019). Such a model would explain the small frequently observed correlations between beta power and RT, as well as associations between “bursty” Mthal activity and beta oscillations in Parkinson Disease (Kühn et al., 2009; Ellens and Leventhal, 2013; Devergnas et al., 2015; Reis et al., 2019). If delta phase-modulated Mthal single unit activity both initiates movement and drives cortical beta oscillations, one would expect weak correlations between beta power and RT. Note that this model does not exclude the possibility that other sources of beta oscillations (e.g., intrinsic basal ganglia oscillators, McCarthy et al., 2011; Tachibana et al., 2011; Mirzaei et al., 2017) are independently associated with behavior.

The identity of directionally- and non-directionally selective units has important implications for understanding subcortical mechanisms of motor control, as well as how LFP oscillations are generated and regulated. One possibility is that these functionally-defined units are anatomically defined by layer-specific cortical projections. Thalamic afferent activity in motor cortical layer 1 is correlated with the speed of individual lever pulls performed by mice, and layer 3 afferents are active at movement initiation (Tanaka et al., 2018). These patterns are strikingly similar to our directionally- and non-directionally selective units, respectively (Gaidica et al., 2018). Furthermore, modeling studies suggest that coordinated layer-specific thalamocortical inputs drive cortical beta oscillations (Sherman et al., 2016). This model requires precisely-timed layer 1 input, which could be provided by directionally-selective units given the correlation between their activity and beta oscillatory power.

A related possibility is that directionally- and non-directionally selective units reside in basal ganglia- and cerebellar-recipient Mthal, respectively. Mthal comprises two mostly non-overlapping subregions defined by basal ganglia or cerebellar afferents (Deniau et al., 1992; Kuramoto et al., 2011) that tend to project to cortical layers 1 and 3/5 respectively (though not with 100% certainty) (Herkenham, 1980; Kuramoto et al., 2009, 2015; Tanaka et al., 2018). In addition to the evidence suggesting that functionally defined Mthal units may have layer-specific projections, several observations also suggest that directionally-selective units reside in basal ganglia-recipient Mthal. First, directionally selective unit activity predicts features of task performance commonly attributed to the basal ganglia (action selective and movement vigor). Indeed basal ganglia manipulations influence RT, MT, and movement direction in nearly identical tasks (Carli et al., 1985; Dowd and Dunnett, 2005; Leventhal et al., 2014). Second, directionally selective units were consistently entrained to delta oscillations during wakefulness, as are basal ganglia-recipient Mthal units (compared to cerebellar-recipient units) under anesthesia (Nakamura et al., 2014). Finally, directionally-selective unit activity predicted beta oscillatory power, which is associated with basal ganglia-thalamocortical circuitry (Leventhal et al., 2012; López-Azcárate et al., 2013; Brittain and Brown, 2014; Feingold et al., 2015).

In summary, we found complex relationships between Mthal LFP oscillations, single unit activity, and performance of a two-alternative forced choice task. These results support a model in which low frequency LFP oscillations either modulate or reflect Mthal neuronal excitability, which in turn drives movement initiation and regulates higher frequency (beta/low gamma) oscillations. These results potentially explain consistently observed correlations between delta phase, beta power, and behavior. A critical open question is the identity of functionally distinct Mthal neuronal populations, which we predict receive distinct subcortical afferents and project to different cortical layers. These predictions have significant implications for understanding subcortical contributions to motor control, and should be testable by combining modern anatomic tracing techniques with high-density electrophysiology and/or optogenetics.

## Materials and Methods

### Data Collection

Detailed data collection methods have been previously described (Gaidica et al., 2018). All animal procedures were approved by the Institutional Animal Care and Use Committee of the University of Michigan. 5 adult male Long-Evans rats (Charles River Laboratories, Wilmington, MA) were housed on a reverse light/dark cycle and food restricted on training days. Operant chambers (ENV-009 Med Associates) were outfitted with 5 illuminated nose ports along one side with an opposite-facing reward port (Figure 1B). Rats were progressively trained to poke one of three illuminated center ports (only one port illuminated per trial) and then, after a variable delay (0.5–1 s, pulled from a uniform distribution), instructed to poke a neighboring port based on a brief low (1 kHz, “go left”) or high (4 kHz, “go right”) pitched tone. Correct trials were rewarded with a 45 mg sucrose pellet at the reward port. Rats were required to perform 80% of trials correctly for three sequential 1-hour sessions before being implanted.

Electrophysiological implants were designed in SolidWorks and printed at the University of Michigan 3D Lab using biocompatible resins. Tetrodes spun from 12 μm wire (Sanvik PX000004) or 50 μm single wire electrodes (California Fine Wire) were interfaced with a Tucker Davis Technologies amplifier system (TDT, ZD64, AC32, PZ4, RZ2, and RS4) using a custom printed circuit board (Advanced Circuits). The entire electrode assembly was driven down with a single precision drive screw. Immediately before surgery, the tetrodes (but not single wires) were gold plated according to a third-party protocol (Neuralynx) and impedances for all electrodes were recorded using a nanoZ (White Matter) impedance tester. All implants were surgically placed with the electrodes residing above the final recording site (Mthal; AP: −3.1 mm, ML: 1.2 mm, DV: -7.1 mm) with a ground and reference screw placed over the cerebellum contacting cerebral spinal fluid. Rats recovered for one week before retraining.

Electrodes were driven roughly 60 μm after each recording day. Wideband (0.1*−*10 kHz) neural signals were recorded at 24 kHz with the TDT system, which was interfaced with custom LabVIEW behavioral software to record behavior timestamps. Single units were sorted in Offline Sorter (Plexon) (Gaidica et al., 2018).

### Data Analysis

All data analysis was performed using MATLAB software which was routinely versioned using Git. The wideband data were decimated by a factor of 16 (from 24 kHz to 1.5 kHz) using the MATLAB *decimate* function. Only correct trials that did not contain wideband artifacts were included in our analysis (n = 2,248 from 30 sessions) using the following exclusion criteria. The signal was converted to a z-score based on the mean and standard deviation from all trials in that session. If the z-score exceeded 5 for more than 5% of the trial length, the trial was excluded. We used the same single unit population (n = 366) from a previous study that did not consider LFP interactions within neuronal firing (Gaidica et al., 2018).

#### Power Spectrum

We visually inspected the raw data from all electrodes from each session and ranked their recording quality to select electrodes with no high amplitude artifacts. This enabled us to use a single, high quality LFP signal from each session for our analyses. In addition, for spike-power and spike-phase correlations, we selected LFP signals from wires where the spikes were not recorded, minimizing the possible influence of the spike waveform itself on the LFP.

We separately analyzed epochs during which the rat was engaged in the task (“in-trial”, between the Cue and Reward) and between trials (“inter-trial”, after the Reward and before the Cue). We created the in-trial power spectrum by concatenating the wideband LFP from all in-trial time periods from a single session. Next, we performed a Fourier transform (*fft* in MATLAB) to obtain the power-frequency spectrum. In order to obtain an average spectrum for all sessions, we divdided the spectrum by the average power of the 70*−*150 Hz segment, which accounted for the variability associated with using different types of electrodes. We present the average spectrum using a conservative (0.2%) smoothing window (*smooth* in MATLAB, Figure 1C). We created the inter-trial power spectrum in the same way but selected inter-trial segments of the LFP equal to the in-trial duration.

#### LFP Correlates of Behavior

A complex scalogram (1–200 Hz, 30 steps log-scale) was computed for each trial by applying a bank of Gabor filters to the raw data (Wallisch et al., 2013). The peri-event (± 1 s) window was extracted from a buffered data series to eliminate filter edge effects. LFP power was calculated by taking the squared magnitude of the complex spectrum. For each session, we determined the mean (μ_baseline_) and standard deviation (*σ*_baseline_) ofbaseline power using a 2 second window leading up to the Cue event for each trial. The average μ_baseline_ and *σ*_baseline_ for each session (μ_session_ and *σ*_session_, respectively) allowed us to z-score the peri-event power of each trial.

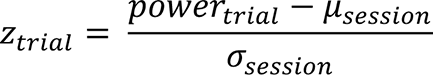

LFP phase was determined using the *angle* function in MATLAB on the complex scalogram. The mean resultant vector length (MRL) for phase data was computed using the *circ_r* function from the Circular Statistics Toolbox (CircStat) for MATLAB (Berens, 2009). Z-scored power and the raw MRL values were calculated for each session and reported as the mean across sessions (Figure 2).

#### Power Comodulation

Power comodugrams were generated using the *corr* function in MATLAB (Pearson’s correlation). For each session, the power from all trials for each event (± 0.5 s) was concatenated and used to calculate the correlation coefficients between the power time series for all frequency pairs (1–200 Hz, 30 steps log-scale). Trial-shuffled comodugrams were generated by pairing the LFP power time series at frequency *f_1_* from one trial with the power time series at frequency *f_2_* from a randomly selected trial from the same session. This calculation was repeated for each *f_1_*-*f_2_* frequency pair 100 times for each session to generate surrogate comodugrams. Actual and surrogate comodugrams are presented as the average over all sessions (Figure 3).

#### Phase-amplitude Coupling (PAC)

We quantified the strength of PAC using established methods (Canolty et al., 2006). A complex scalogram (1–200 Hz, 30 steps log-scale) was computed for a peri-event time window (± 0.5s) for each trial. For each session, we concatenated data from all correct trials for each event. Thus, we obtained a complex time series for each event that was *n*-seconds long, where *n* is the number of trials in a session. We then obtained the time-series phase (*Φ(t)*) by applying the *angle* function in MATLAB and amplitude (*A(t)*) by taking the squared magnitude. These data were used to determine the PAC between pairs of frequencies (*m*,*n*) across all events, with the constraint that the amplitude frequency *m* was always greater than or equal to the phase frequency *n*. We achieved this by first creating a composite phase-amplitude signal (*z_t_*) from the session-wide time series data:

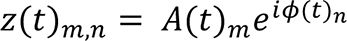

The mean (*M_m,n_*) of *z(t)_m,n_* quantifies the deviation of *z(t)_m,n_* from a radially symmetric distribution of high frequency LFP amplitudes across low frequency phases. To account for the possibility that *Φ(t)_n_* is not uniformly distributed, we normalized *M_m,n_* for each session using 200 surrogates generated by adding a random lag *τ* to A(t).

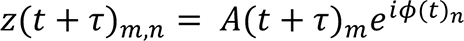

*M_surr_* is the mean of *z(t+τ)_m,n_* and is calculated separately for each surrogate phase-amplitude analysis. The mean (μ_Msurr_) and standard deviation (*σ*_Msurr_) of the surrogate distribution were calculated using *normfit* in MATLAB (where the input was all 200 *M_surr_* values). We report the modulation index (*MI_m,n_*) as the magnitude of the normalized *M_m,n_* (Figure 4).

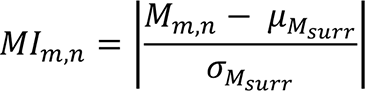

A p-value was obtained for each phase-amplitude pair in the *MI* matrix using *normcdf* in MATLAB with the ‘upper’ option to compute right-tailed probabilities. We corrected for multiple comparisons (Bonferroni method) by multiplying the p-values by the number of elements in *MI_m,n_* (N = 30 *×* 30). For example, using *α* = 0.05, the z-score contained in *MI* must exceed 3.87 to reach significance (determined using the *norminv* function in MATLAB on *α* ÷ N).

To determine if PAC was present independent of correlations between LFP features and behavior, we recalculated surrogate MIs 1,000 times from a composite signal where the trial order of *A(t)* was shuffled (Stark and Abeles, 2005). This allowed us to generate a statistical measure for the fraction of shuffled *MI*s greater or less than the true *MI*.

#### Single Unit Entrainment

We extracted the instantaneous phase of the LFP from the complex spectrum (using the MATLAB *angle* function) for each spike timestamp within equal duration in-trial and inter-trial periods. Next, we performed a Rayleigh test for non-uniformity of circular data (CircStat *circ_rtest* function) (Berens, 2009) for the compiled phases to obtain a p-value to reject the null hypothesis that spike timing is uniformly distributed from −180° to 180° (Figure 5A). To determine if the number of units significantly entrained (p < 0.05) to each frequency was greater than chance, we generated firing rate matched, Poisson distributed spike trains for each unit and recalculated the p-values 1,000 times. We used the same data to calculate the mean MRL of LFP phase at each spike timestamp for each unit population (Figure 5B), and similarly compared it against Poisson spikes. P-values were determined as the fraction of significantly entrained unit percentages/MRL values from surrogate calculations that were greater than the actual value.

To determine if the preferred firing phase was consistent across units, we generated spike histograms for each unit across 12 linearly-spaced phase bins between −180° and 180° (Figure 6).Each unit histogram was normalized by dividing each bin count by the total number of spikes for that unit to account for spike rate. We used the same method described above to generate surrogate Poisson spike-phase histograms, which were used to assess the significance of single unit phase preferences.

#### Spike-power Cross Correlations

We used a cross correlation to determine the relationship between LFP power and single unit activity for equal duration in-trial and inter-trial periods. First, we generated a session-wide continuous spike density estimate (SDE) for each unit and trial by convolving the vector of discrete spiking events with a 50 ms Gaussian kernel (Wallisch et al., 2013). Next, we extracted the relevant SDE segments for the in-trial and inter-trial periods. We cross correlated these data with LFP power (1–200 Hz, 30 steps log-scale) on a per-trial basis using the *xcorr* function in MATLAB with the ‘coeff’ option so that the autocorrelations at zero lag equal 1. Cross correlation matrices are presented as the mean over all trials and sessions (Figure 7). We recalculated each cross correlation using a firing rate matched, Poisson distributed spike train 20 times, giving us a distribution of correlation values across time for each frequency. The maximum and minimum of that distribution are where we considered values to be significantly different from chance (Figure 8).

#### LFP Correlates of Performance

To determine relationships between LFP features, RT, and MT, we used peri-event (± 1 s) power and phase data for each frequency (1–200 Hz, 30 steps log-scale) and all trials. For each time point and frequency, we created a 1-by-*n* array of power (or phase) values, where *n* was the number of trials in that session, along with a 1-by-*n* array of the RT (or MT) values for each trial. We used these two arrays as inputs to the *corr* function in MATLAB to calculate Spearman’s correlation coefficient for power-RT/MT, and the *circ_corrcl* function (CircStat toolbox, Berens, 2009) for phase-RT/MT correlations. Therefore, each time-frequency pair generated a single correlation coefficient and associated p-value between power/phase and RT/MT (Figure 9).

---

**Figure 10—figure supplement 1.**
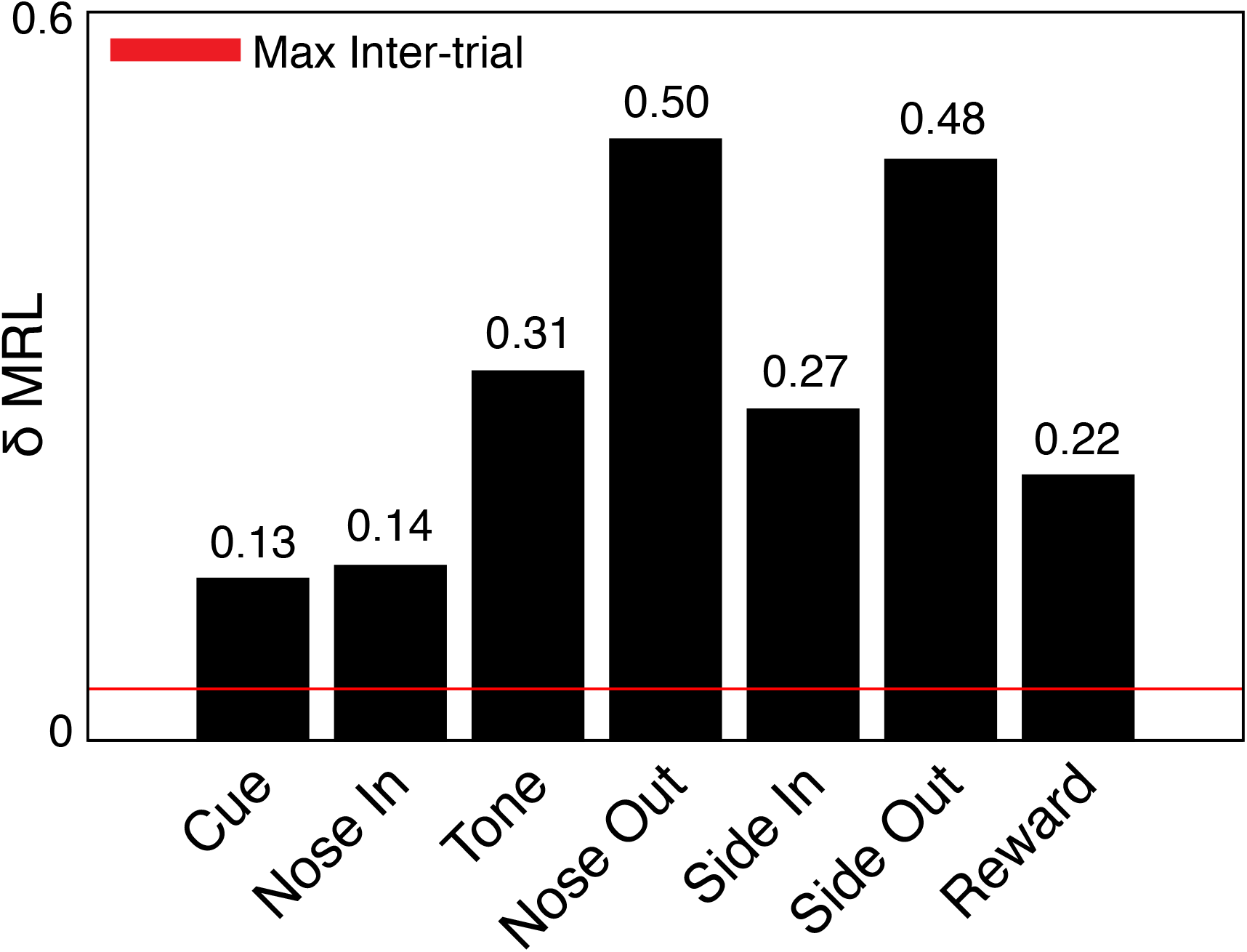
Delta phase becomes aligned to task events before Nose Out. The mean resultant length (MRL) for delta phase (2.5 Hz) at the time of each event. The red line indicates the maximum inter-trial MRL obtained by resampling delta phase during the inter-trial period (see Materials and Methods).

## Source Data Files

**Figure 2 – Source Data 1.** The data structures in the file *Figure2_SourceData.mat* contain the time-series power and mean resultant length (phase) calculations for all sessions. The accompanying m-file *Figure2_SourceCode.m* describes the data structures and produces the grand average scalograms and MRL heatmaps using the package contained in https://github.com/LeventhalLab/eLifeMthal.

**Figure 3 – Source Data 1.** The data structures in the file *Figure3_SourceData.mat* contain the frequency-frequency comodugram data for all sessions. The data structures include a shuffled condition, as well data for the inter-trial interval. The accompanying m-file *Figure3_SourceCode.m* describes the data structures and produces the grand average frequency-frequency comodugrams using the package contained in https://github.com/LeventhalLab/eLifeMthal.

**Figure 4 – Source Data 1.** The data structures in the file *Figure4_SourceData.mat* contain phase-amplitude coupling data for all sessions as frequency-frequency matrices. Values were stored as modulation index (MI, see Materials and Methods). The data structures include a shuffled condition, data for the inter-trial interval, and a separate matrix for calculated p-values. The accompanying m-file *Figure4_SourceCode.m* describes the data structures and produces the grand average frequency-frequency MI matrices and circles areas where p < 0.05 in red using the package contained in https://github.com/LeventhalLab/eLifeMthal.

**Figure 5 – Source Data 1.** The data structures in the file *Figure5_SourceData.mat* contain entrainment data for each unit (n = 366), specifically, the p-value associated with a Rayleigh test for non-uniformity across all frequencies. Unit indices for directionally selective and non-directionally selective units are stored in arrays. Entrainment data from Poisson spike distributions to indicate chance levels of unit-phase correlations (see Materials and Methods) are save alongside the real data. The accompanying m-file *Figure5_SourceCode.m* describes the data structures and displays the fraction of each unit group where p < 0.05 using the package contained in https://github.com/LeventhalLab/eLifeMthal.

**Figure 6 – Source Data 1.** The data structures in the file *Figure6_SourceData.mat* contain entrainment data for each unit (n = 366) including normalized spike-phase histograms for real and Poisson spike trains. The accompanying m-file *Figure6_SourceCode.m* describes the data structures and displays entrainment histograms for unit types for the in-trial and inter-trial time periods for the delta band (2.5 Hz) using the package contained in https://github.com/LeventhalLab/eLifeMthal. Spike preference plots are grand averages of the spike-phase histograms.

**Figure 9 – Source Data 1.**The data structures in the file *Figure9_SourceData.mat* contain the reaction time and movement time correlations across all frequencies for power and phase and all sessions (see Materials and Methods for calculation methods). P-values for each power-phase-frequency correlation was saved for each session. The accompanying m-file *Figure9_SourceCode.m* describes the data structures and displays power and phase correlations as a heatmap and overlays the number of sessions where the correlation was significant (p < 0.05) for delta (2.5 Hz) phase and beta (20 Hz) power using the package contained in https://github.com/LeventhalLab/eLifeMthal.

## References

Anastassiou CA, Montgomery SM, Barahona M, Buzsáki G, Koch C (2010) The effect of spatially inhomogeneous extracellular electric fields on neurons. J Neurosci, 30:1925–1936.

Armstrong S, Sale MV, Cunnington R (2018) Neural Oscillations and the Initiation of Voluntary Movement. Front Psychol, 9:2509.

Arnal LH, Doelling KB, Poeppel D (2015) Delta-Beta Coupled Oscillations Underlie Temporal Prediction Accuracy. Cereb Cortex, 25:3077–3085.

Baker SN, Olivier E, Lemon RN (1997) Coherent oscillations in monkey motor cortex and hand muscle EMG show task-dependent modulation. J Physiol, 501:225–241.

Bansal AK, Vargas-Irwin CE, Truccolo W, Donoghue JP (2011) Relationships among low-frequency local field potentials, spiking activity, and three-dimensional reach and grasp kinematics in primary motor and ventral premotor cortices. J Neurophysiol, 105:1603–1619.

Bastos AM, Briggs F, Alitto HJ, Mangun GR, Usrey WM (2014) Simultaneous recordings from the primary visual cortex and lateral geniculate nucleus reveal rhythmic interactions and a cortical source for γ-band oscillations. J Neurosci, 34:7639–7644.

Belluscio MA, Mizuseki K, Schmidt R, Kempter R, Buzsáki G (2012) Cross-Frequency Phase-Phase Coupling between Theta and Gamma Oscillations in the Hippocampus. J Neurosci, 32:423–435.

Berens P (2009) CircStat: a MATLAB toolbox for circular statistics. J Stat Softw, 31:1–21.

Berke JD, Okatan M, Skurski J, Eichenbaum HB (2004) Oscillatory entrainment of striatal neurons in freely moving rats. Neuron, 43:883–896.

Breska A, Deouell LY (2017) Neural mechanisms of rhythm-based temporal prediction: Delta phase-locking reflects temporal predictability but not rhythmic entrainment. PLoS Biol, 15:e2001665.

Brittain J-SS, Brown P (2014) Oscillations and the basal ganglia: motor control and beyond. Neuroimage, 85 Pt 2:637–647.

Brown P (2006) Bad oscillations in Parkinson’s disease. J Neural Transm Suppl, 27–30.

Canolty RT, Edwards E, Dalal SS, Soltani M, Nagarajan SS, Kirsch HE, Berger MS, Barbaro NM, Knight RT (2006) High gamma power is phase-locked to theta oscillations in human neocortex. Science, 313:1626–1628.

Canolty RT, Soltani M, Dalal SS, Edwards E, Dronkers NF, Nagarajan SS, Kirsch HE, Barbaro NM, Knight RT (2007) Spatiotemporal dynamics of word processing in the human brain. Front Neurosci, 1:185–196.

Carli M, Evenden JL, Robbins TW (1985) Depletion of unilateral striatal dopamine impairs initiation of contralateral actions and not sensory attention. Nature, 313:679–682.

Cohen MX, Elger CE, Fell J (2009) Oscillatory activity and phase-amplitude coupling in the human medial frontal cortex during decision making. J Cogn Neurosci, 21:390–402.

Crunelli V, David F, Lőrincz ML, Hughes SW (2015) The thalamocortical network as a single slow wave-generating unit. Curr Opin Neurobiol, 31:72–80.

Crunelli V, Lőrincz ML, Connelly WM, David F, Hughes SW, Lambert RC, Leresche N, Errington AC (2018) Dual function of thalamic low-vigilance state oscillations: rhythm-regulation and plasticity. Nat Rev Neurosci, 19:107–118.

de Cheveigné A, Nelken I (2019) Filters: When, Why, and How (Not) to Use Them. Neuron, 102:280–293.

de Hemptinne C, Ryapolova-Webb ES, Air EL, Garcia PA, Miller KJ, Ojemann JG, Ostrem JL, Galifianakis NB, Starr PA (2013) Exaggerated phase-amplitude coupling in the primary motor cortex in Parkinson disease. Proc Natl Acad Sci U S A, 110:4780–4785.

Dejean C, Arbuthnott G, Wickens JR, Le Moine C, Boraud T, Hyland BI (2011) Power Fluctuations in Beta and Gamma Frequencies in Rat Globus Pallidus: Association with Specific Phases of Slow Oscillations and Differential Modulation by Dopamine D1 and D2 Receptors. J Neurosci, 31:6098–6107.

Deniau JM, Kita H, Kitai ST (1992) Patterns of termination of cerebellar and basal ganglia efferents in the rat thalamus. Strictly segregated and partly overlapping projections. Neurosci Lett, 144:202–206.

Devergnas A, Chen E, Ma Y, Hamada I, Pittard D, Kammermeier S, Mullin AP, Faundez V, Lindsley CW, Jones C, Smith Y, Wichmann T (2015) Anatomical Localization of CaV3.1 Calcium Channels and Electrophysiological Effects of T-type Calcium Channel Blockade in the Thalamus of MPTP-Treated Monkeys. J Neurophysiol, jn.00858.2015.

Donoghue JP, Sanes JN, Hatsopoulos NG, Gaál G (1998) Neural discharge and local field potential oscillations in primate motor cortex during voluntary movements. J Neurophysiol, 79:159–173.

Dowd E, Dunnett SB (2005) Comparison of 6-hydroxydopamine-induced medial forebrain bundle and nigrostriatal terminal lesions in a lateralised nose-poking task in rats. Behav Brain Res, 159:153–161.

Ellens DJ, Leventhal DK (2013) Review: electrophysiology of basal ganglia and cortex in models of Parkinson disease. J Parkinsons Dis, 3:241–254.

Engel AK, Fries P (2010) Beta-band oscillations--signalling the status quo? Curr Opin Neurobiol, 20:156–165.

Feingold J, Gibson DJ, DePasquale B, Graybiel AM (2015) Bursts of beta oscillation differentiate postperformance activity in the striatum and motor cortex of monkeys performing movement tasks. Proc Natl Acad Sci U S A, 112:13687–13692.

Fiebelkorn IC, Snyder AC, Mercier MR, Butler JS, Molholm S, Foxe JJ (2013) Cortical cross-frequency coupling predicts perceptual outcomes. Neuroimage, 69:126–137.

Fogerson PM, Huguenard JR (2016) Tapping the Brakes: Cellular and Synaptic Mechanisms that Regulate Thalamic Oscillations. Neuron, 92:687–704.

Fries P (2015) Rhythms for Cognition: Communication through Coherence. Neuron, 88:220–235.

Friston KJ, Bastos AM, Pinotsis D, Litvak V (2015) LFP and oscillations-what do they tell us. Curr Opin Neurobiol, 31:1–6.

Fujisawa S, Buzsáki G (2011) A 4 Hz oscillation adaptively synchronizes prefrontal, VTA, and hippocampal activities. Neuron, 72:153–165.

Gaidica M, Hurst A, Cyr C, Leventhal DK (2018) Distinct Populations of Motor Thalamic Neurons Encode Action Initiation, Action Selection, and Movement Vigor. J Neurosci, 38:6563–6573.

Gilbertson T, Lalo E, Doyle L, Di Lazzaro V, Cioni B, Brown P (2005) Existing motor state is favored at the expense of new movement during 13-35 Hz oscillatory synchrony in the human corticospinal system. J Neurosci, 25:7771–7779.

Grabot L, Kononowicz TW, Dupré la Tour T, Gramfort A, Doyère V, van Wassenhove V (2019) The strength of alpha-beta oscillatory coupling predicts motor timing precision. J Neurosci,

Halassa MM, Siegle JH, Ritt JT, Ting JT, Feng G, Moore CI (2011) Selective optical drive of thalamic reticular nucleus generates thalamic bursts and cortical spindles. Nat Neurosci, 14:1118–1120.

Hamel-Thibault A, Thénault F, Whittingstall K, Bernier PM (2018) Delta-Band Oscillations in Motor Regions Predict Hand Selection for Reaching. Cereb Cortex, 28:574–584.

Herkenham M (1980) Laminar organization of thalamic projections to the rat neocortex. Science, 207:532–535.

Igarashi J, Isomura Y, Arai K, Harukuni R, Fukai T (2013) A $λ$-$γ$ Oscillation Code for Neuronal Coordination during Motor Behavior. J Neurosci, 33:18515–18530.

Jones SR, Pritchett DL, Sikora MA, Stufflebeam SM, Hämäläinen M, Moore CI (2009) Quantitative analysis and biophysically realistic neural modeling of the MEG mu rhythm: rhythmogenesis and modulation of sensory-evoked responses. J Neurophysiol, 102:3554–3572.

Khanna P, Carmena JM (2017) Beta band oscillations in motor cortex reflect neural population signals that delay movement onset. Elife, 6

Kühn AA, Tsui A, Aziz T, Ray N, Brücke C, Kupsch A, Schneider G-HH, Brown P (2009) Pathological synchronisation in the subthalamic nucleus of patients with Parkinson’s disease relates to both bradykinesia and rigidity. Exp Neurol, 215:380–387.

Kuramoto E, Fujiyama F, Nakamura KC, Tanaka Y, Hioki H, Kaneko T (2011) Complementary distribution of glutamatergic cerebellar and GABAergic basal ganglia afferents to the rat motor thalamic nuclei. Eur J Neurosci, 33:95–109.

Kuramoto E, Furuta T, Nakamura KC, Unzai T, Hioki H, Kaneko T (2009) Two types of thalamocortical projections from the motor thalamic nuclei of the rat: a single neuron-tracing study using viral vectors. Cereb Cortex, 19:2065–2077.

Kuramoto E, Ohno S, Furuta T, Unzai T, Tanaka YR, Hioki H, Kaneko T (2015) Ventral medial nucleus neurons send thalamocortical afferents more widely and more preferentially to layer 1 than neurons of the ventral anterior-ventral lateral nuclear complex in the rat. Cereb Cortex, 25:221–235.

Lakatos P, Karmos G, Mehta AD, Ulbert I, Schroeder CE (2008) Entrainment of neuronal oscillations as a mechanism of attentional selection. Science, 320:110–113.

Lakatos P, Shah AS, Knuth KH, Ulbert I, Karmos G, Schroeder CE (2005) An oscillatory hierarchy controlling neuronal excitability and stimulus processing in the auditory cortex. J Neurophysiol, 94:1904–1911.

Leventhal DK, Gage GJ, Schmidt R, Pettibone JR, Case AC, Berke JD (2012) Basal Ganglia Beta oscillations accompany cue utilization. Neuron, 73:523–536.

Leventhal DK, Stoetzner CR, Abraham R, Pettibone J, DeMarco K, Berke JD (2014) Dissociable effects of dopamine on learning and performance within sensorimotor striatum. Basal Ganglia, 4:43–54.

Lofredi R, Tan H, Neumann W-J, Yeh C-H, Schneider G-H, Kühn AA, Brown P (2019) Beta bursts during continuous movements accompany the velocity decrement in Parkinson’s disease patients. Neurobiology of Disease,

López-Azcárate J, Nicolás MJ, Cordon I, Alegre M, Valencia M, Artieda J (2013) Delta-mediated cross-frequency coupling organizes oscillatory activity across the rat cortico-basal ganglia network. Front Neural Circuits, 7:155.

Mak-McCully RA, Rolland M, Sargsyan A, Gonzalez C, Magnin M, Chauvel P, Rey M, Bastuji H, Halgren E (2017) Coordination of cortical and thalamic activity during non-REM sleep in humans. Nat Commun, 8:15499.

Mallet N, Pogosyan A, Márton LF, Bolam JP, Brown P, Magill PJ (2008) Parkinsonian beta oscillations in the external globus pallidus and their relationship with subthalamic nucleus activity. J Neurosci, 28:14245–14258.

Manning JR, Jacobs J, Fried I, Kahana MJ (2009) Broadband shifts in local field potential power spectra are correlated with single-neuron spiking in humans. J Neurosci, 29:13613–13620.

Masimore B, Schmitzer-Torbert NC, Kakalios J, Redish AD (2005) Transient striatal gamma local field potentials signal movement initiation in rats. Neuroreport, 16:2021–2024.

McAfee SS, Liu Y, Dhamala M, Heck DH (2018) Thalamocortical Communication in the Awake Mouse Visual System Involves Phase Synchronization and Rhythmic Spike Synchrony at High Gamma Frequencies. Front Neurosci, 12:837.

McCarthy MM, Moore-Kochlacs C, Gu X, Boyden ES, Han X, Kopell N (2011) Striatal origin of the pathologic beta oscillations in Parkinson’s disease. Proc Natl Acad Sci U S A,

Meidahl AC, Moll CKE, van Wijk B, Gulberti A, Tinkhauser G, Westphal M, Engel AK, Hamel W, Brown P, Sharott A (2019) Synchronised spiking activity underlies phase amplitude coupling in the subthalamic nucleus of Parkinson’s disease patients. Neurobiol Dis,

Mirzaei A, Kumar A, Leventhal D, Mallet N, Aertsen A, Berke J, Schmidt R (2017) Sensorimotor Processing in the Basal Ganglia Leads to Transient Beta Oscillations during Behavior. 37:11220–11232.

Murthy VN, Fetz EE (1992) Coherent 25-to 35-Hz oscillations in the sensorimotor cortex of awake behaving monkeys. Proc Natl Acad Sci U S A, 89:5670–5674.

Nakamura KC, Sharott A, Magill PJ (2014) Temporal coupling with cortex distinguishes spontaneous neuronal activities in identified basal ganglia-recipient and cerebellar-recipient zones of the motor thalamus. Cereb Cortex, 24:81–97.

Neske GT (2015) The Slow Oscillation in Cortical and Thalamic Networks: Mechanisms and Functions. Front Neural Circuits, 9:88.

Pesaran B, Vinck M, Einevoll GT, Sirota A, Fries P, Siegel M, Truccolo W, Schroeder CE, Srinivasan R (2018) Investigating large-scale brain dynamics using field potential recordings: analysis and interpretation. Nat Neurosci, 21:903–919.

Pfurtscheller G, Stancák A, Neuper C (1996) Post-movement beta synchronization. A correlate of an idling motor area? Electroencephalogr Clin Neurophysiol, 98:281–293.

Pogosyan A, Gaynor LD, Eusebio A, Brown P (2009) Boosting cortical activity at Beta-band frequencies slows movement in humans. Curr Biol, 19:1637–1641.

Ray S, Crone NE, Niebur E, Franaszczuk PJ, Hsiao SS (2008) Neural correlates of high-gamma oscillations (60-200 Hz) in macaque local field potentials and their potential implications in electrocorticography. J Neurosci, 28:11526–11536.

Reis C, Sharott A, Magill PJ, van Wijk BCM, Parr T, Zeidman P, Friston KJ, Cagnan H (2019) Thalamocortical dynamics underlying spontaneous transitions in beta power in Parkinsonism. Neuroimage, 193:103–114.

Rule ME, Vargas-Irwin C, Donoghue JP, Truccolo W (2018) Phase reorganization leads to transient β-LFP spatial wave patterns in motor cortex during steady-state movement preparation. J Neurophysiol, 119:2212–2228.

Saalmann YB, Pinsk MA, Wang L, Li X, Kastner S (2012) The pulvinar regulates information transmission between cortical areas based on attention demands. Science, 337:753–756.

Saleh M, Reimer J, Penn R, Ojakangas CL, Hatsopoulos NG (2010) Fast and slow oscillations in human primary motor cortex predict oncoming behaviorally relevant cues. Neuron, 65:461–471.

Schmidt R, Leventhal DK, Mallet N, Chen F, Berke JD (2013) Canceling actions involves a race between basal ganglia pathways. Nat Neurosci, 16:1118–1124.

Schroeder CE, Lakatos P (2009) Low-frequency neuronal oscillations as instruments of sensory selection. Trends Neurosci, 32:9–18.

Sherman MA, Lee S, Law R, Haegens S, Thorn CA, Hämäläinen MS, Moore CI, Jones SR (2016) Neural mechanisms of transient neocortical beta rhythms: Converging evidence from humans, computational modeling, monkeys, and mice. Proceedings of the National Academy of Sciences, 113:E4885–E4894.

Shin H, Law R, Tsutsui S, Moore CI, Jones SR (2017) The rate of transient beta frequency events predicts behavior across tasks and species. Elife, 6

Stark E, Abeles M (2005) Applying resampling methods to neurophysiological data. J Neurosci Methods, 145:133–144.

Stefanics G, Hangya B, Hernádi I, Winkler I, Lakatos P, Ulbert I (2010) Phase entrainment of human delta oscillations can mediate the effects of expectation on reaction speed. J Neurosci, 30:13578–13585.

Tachibana Y, Iwamuro H, Kita H, Takada M, Nambu A (2011) Subthalamo-pallidal interactions underlying parkinsonian neuronal oscillations in the primate basal ganglia. Eur J Neurosci, 34:1470–1484.

Tan H, Debarros J, He S, Pogosyan A, Aziz TZ, Huang Y, Wang S, Timmermann L, Visser-Vandewalle V, Pedrosa DJ, Green AL, Brown P (2019) Decoding voluntary movements and postural tremor based on thalamic LFPs as a basis for closed-loop stimulation for essential tremor. Brain Stimul,

Tanaka YH, Tanaka YR, Kondo M, Terada SI, Kawaguchi Y, Matsuzaki M (2018) Thalamocortical Axonal Activity in Motor Cortex Exhibits Layer-Specific Dynamics during Motor Learning. Neuron, 100:244–258.e12.

Tiganj Z, Chevallier S, Monacelli E (2014) Influence of extracellular oscillations on neural communication: a computational perspective. Front Comput Neurosci, 8:9.

Torrecillos F, Tinkhauser G, Fischer P, Green AL, Aziz TZ, Foltynie T, Limousin P, Zrinzo L, Ashkan K, Brown P, Tan H (2018) Modulation of Beta Bursts in the Subthalamic Nucleus Predicts Motor Performance. J Neurosci, 38:8905–8917.

Tort ABL, Kramer MA, Thorn C, Gibson DJ, Kubota Y, Graybiel AM, Kopell NJ (2008) Dynamic cross-frequency couplings of local field potential oscillations in rat striatum and hippocampus during performance of a T-maze task. Proc Natl Acad Sci U S A, 105:20517–20522.

van Wijk BCM (2017) Is Broadband Gamma Activity Pathologically Synchronized to the Beta Rhythm in Parkinson’s Disease. J Neurosci, 37:9347–9349.

Wallisch P, Lusignan ME, Benayoun MD, Baker TI, Dickey AS, Hatsopoulos NG (2013) MATLAB for Neuroscientists: An Introduction to Scientific Computing in MATLAB. Academic Press.

Wang P, Göschl F, Friese U, König P, Engel AK (2019) Long-range functional coupling predicts performance: Oscillatory EEG networks in multisensory processing. Neuroimage, 196:114–125.

Watson BO, Ding M, Buzsáki G (2018) Temporal coupling of field potentials and action potentials in the neocortex. Eur J Neurosci, 48:2482–2497.

Zoefel B, Archer-Boyd A, Davis MH (2018) Phase entrainment of brain oscillations causally modulates neural responses to intelligible speech. Current Biology, 28:401–408. e5.

